# Antigenic diversity in malaria parasites is maintained on extrachromosomal DNA

**DOI:** 10.1101/2023.02.02.526885

**Authors:** Emily R. Ebel, Bernard Y. Kim, Marina McDew-White, Elizabeth S. Egan, Timothy J.C. Anderson, Dmitri A. Petrov

## Abstract

Sequence variation among antigenic *var* genes enables *Plasmodium falciparum* malaria parasites to evade host immunity. Using long sequence reads from haploid clones from a mutation accumulation experiment, we detect *var* diversity inconsistent with simple chromosomal inheritance. We discover putatively circular DNA that is strongly enriched for *var* genes, which exist in multiple alleles per locus separated by recombination and indel events. Extrachromosomal DNA likely contributes to rapid antigenic diversification in *P. falciparum*.

---

Malaria caused by the parasite *Plasmodium falciparum* is a leading cause of death and disease in tropical regions of the world^1^. Adaptive immunity to malaria is limited, even after repeated infection, by extensive variation in *P. falciparum* antigenic gene families^2–4^. In particular, *var* genes encode PfEMP1 proteins that are exported to the surface of infected red blood cells, where they mediate pathogenic cytoadherence to host endothelial receptors and elicit variant-specific immunity^2^. Each parasite genome contains ∼60 *var* genes distributed among 26 subtelomeric and 9 internal loci. *Var* genes are named after variation in their antigenic properties^5^, driven by extreme amino acid divergence^6^ relative to the rest of the genome. For example, pairs of parasites sampled from the same population share almost no *var* genes with ≥96% sequence identity^7^. However, *var* genes from unrelated parasites share small blocks of homology^6,8^ consistent with a history of recombination or gene conversion among alleles. Recent studies have reported frequent *var* recombination during asexual, mitotic reproduction, which may create millions of new alleles during blood-stage infection^9^.

Our current understanding of *var* diversification relies primarily on short-read sequencing that may yield only a partial picture of *var* genetic diversity. The primary hurdle is structural variation among *var* genes^6,10^, such as copy number variation, which makes it difficult to accurately align short reads to reference genomes. To achieve a more complete understanding of *var* diversity and mutational mechanisms, we generated long sequence reads from a mutation accumulation (MA) experiment in *P. falciparum*^11^ (**Fig 1A**). Specifically, 31 MA lines (MAL) were independently cloned from an isogenic population of the 3D7 reference strain (the ‘Ancestor’) and propagated for 6-12 months (∼90-180 cell divisions). Each MAL was re-cloned to a single cell every 21±4 days (∼10 cell divisions) to minimize selection and allow fixation of *de novo* mutations. Previous Illumina analysis of the core genome of 31 MAL identified an average of 0.55 SNP and 3.42 small indel mutations per MAL over the course of the MA experiment^11^.

**Figure 1.**
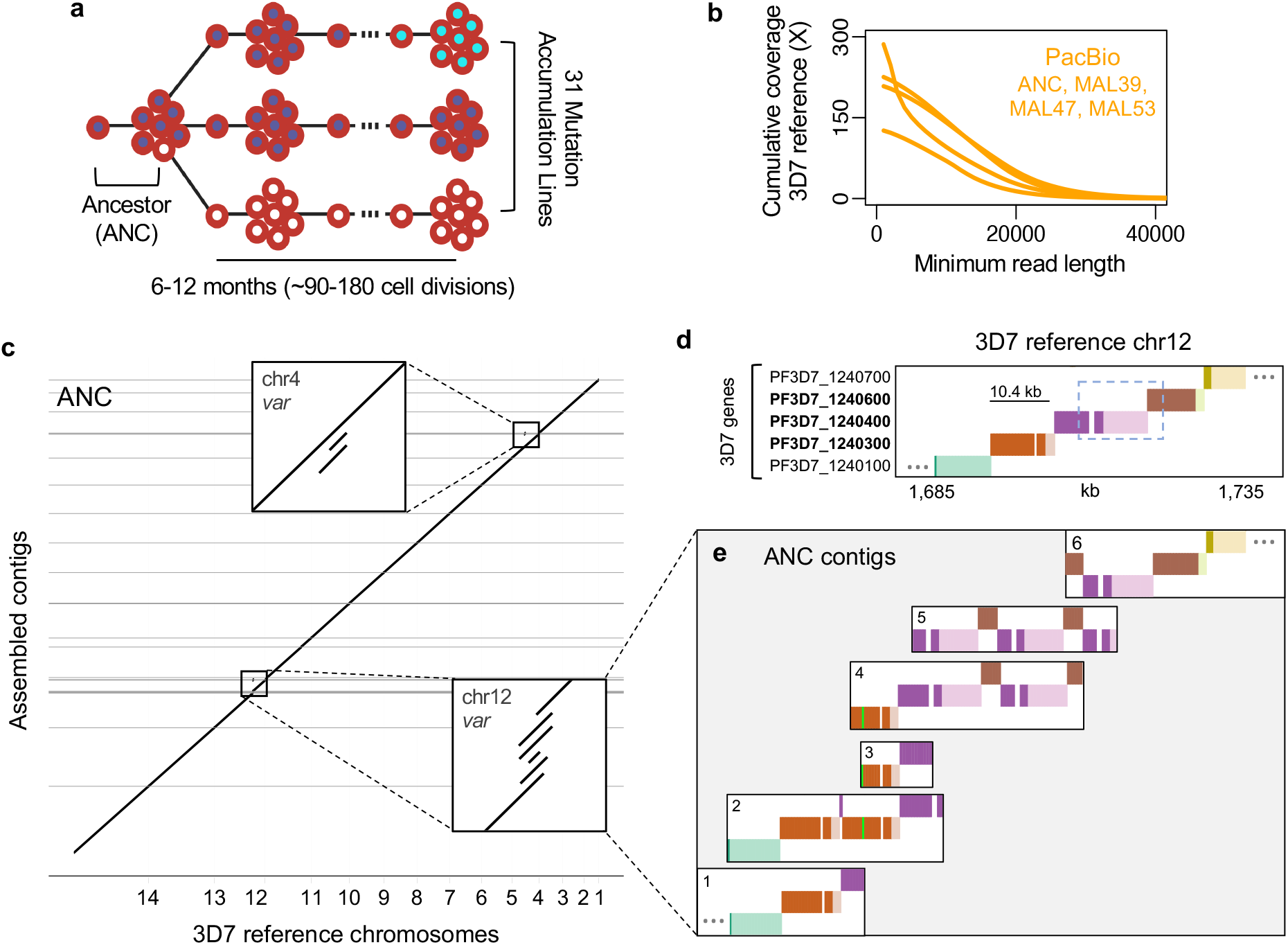
Unexpected clonal polymorphism in PacBio assemblies. **a**, Mutation accumulation experiment with repeated cloning. **b**, PacBio read lengths. **c**, Genome-wide dot plot comparing ANC assembly with 3D7 reference. Insets are internal *var* loci with >1 contig. **d**, Visualization of homology between reference genes (dark colors) and intergenic regions (light colors) on 3D7 chr12 (top) and ANC contigs (bottom). *Var* gene names are bolded. The dashed box outlines a region duplicated on contigs 4-6. The lime green band represents 241 bp from PF3D7_0700100. Ellipses indicate continued sequence (other genes not shown).

To detect large structural mutations at *var* loci, we initially used long PacBio reads (>16kb, **Fig 1B**) to build *de novo* genome assemblies for the Ancestor (ANC) and three MAL (MAL39, MAL47, MAL53). Each of these high-quality assemblies contained few gaps (0-4) and covered ≥99.5% of reference bases with ≥94.4% identity (ANC, **Fig 1C**; others, **EDF 1A**; **Supplementary Table 1**). Nonetheless, we observed that 27 genomic regions were represented by multiple contigs in at least one assembly (**Fig 1C**; EDF 1AB), which could represent structural variation. These incompletely-resolved regions were highly enriched for *var* genes, which comprise 9% of the genome but 69% of unresolved regions (p<0.0001, χ2=228.5). To examine the structural layout of *var* genes on each assembled contig, we developed a Shiny app that draws BLAST homology between contigs and reference genes. As an example, consider an internal *var* locus on chr12, which was identified on multiple contigs in 3 of 4 assemblies (**Fig 1C, EDF 1A**). When we applied our app to this locus in the reference genome, it drew a diagonal line of five sequential genes separated by color and vertical position (**Fig 1D**).

In contrast, when we applied the app to ANC contigs with homology to this locus, it displayed deviations from the reference diagonal that indicate structural variation (**Fig 1DE**). For example, ANC contig 1 matched the reference while contig 2 contained a second copy of *var* PF3D7_1240300 (**Fig 1DE,** orange). This second copy, which was also partially present on contigs 3 and 4, was differentiated from the first copy by two sequence tracts of a few hundred base pairs: one homologous to an adjacent *var* (**Fig 1DE,** purple) and one homologous to a subtelomeric *var* on another chromosome (**Fig 1DE,** lime green). On contigs 4-6, tandem duplications appeared to produce novel, chimeric *var* genes that were in-frame for translation and retained the upstream regulatory sequence of one parent (**Fig 1D,** dashed box; **Fig 1E,** purple/brown). These tandem duplication and recombination events are consistent with known mechanisms of *var* mutation^9^, and we successfully confirmed specific breakpoints using PCR (**EDF 2A**). We also observed similar patterns of structural variation between contigs mapping to two internal *var* loci on chr4 (**EDF 1C-F**). Apparent polymorphism between contigs (1/2, **Fig 1E**; 7/8, **EDF 1D**; 10/11, **EDF 1F**) was unexpected in haploid clones and could reflect assembly errors or *var* polymorphism, including paralog divergence after *var* locus duplication.

To achieve greater resolution of *var* genetic diversity and extend the analysis to additional clones, we generated ultra-long Oxford Nanopore reads from ANC and 16 MAL (**Fig 2A**; mean coverage 11X in reads >100kb). Using the Shiny app, we examined gene structure on individual Nanopore reads mapping to each *var* locus. At the chr12 locus, we observed extensive variation across reads in the copy numbers of *var* PF3D7_1240300 and the novel *var* chimera (**Fig 2B, Fig 1DE,** orange and purple/brown). Most Nanopore reads were not long enough to span both sets of duplications at this locus while being anchored in unique sequence on either end. We therefore divided the locus into two regions for genotyping: one containing the orange *var*, anchored by the adjacent teal and purple genes; and one containing the purple and brown *var*, anchored by the adjacent orange and dark yellow genes (see **Fig 1D** for reference gene structure). Using only reads anchored by this definition, we identified 6 distinct orange alleles and 7 purple/brown alleles across ANC and the 16 MAL (**Fig 2B, A-F, Z-T**). Variation across Nanopore alleles reflected a history of indel and recombination events that altered *var* structure and copy number, consistent with the PacBio contigs (**Fig 1E**). Individual reads additionally revealed rarer variation, such as short tracts of recombination between adjacent *var* genes (**Fig 2B** allele U). We observed similarly high levels of structural variation across reads from three other internal *var* loci on chr7 and chr4 (**EDF 3**).

**Figure 2.**
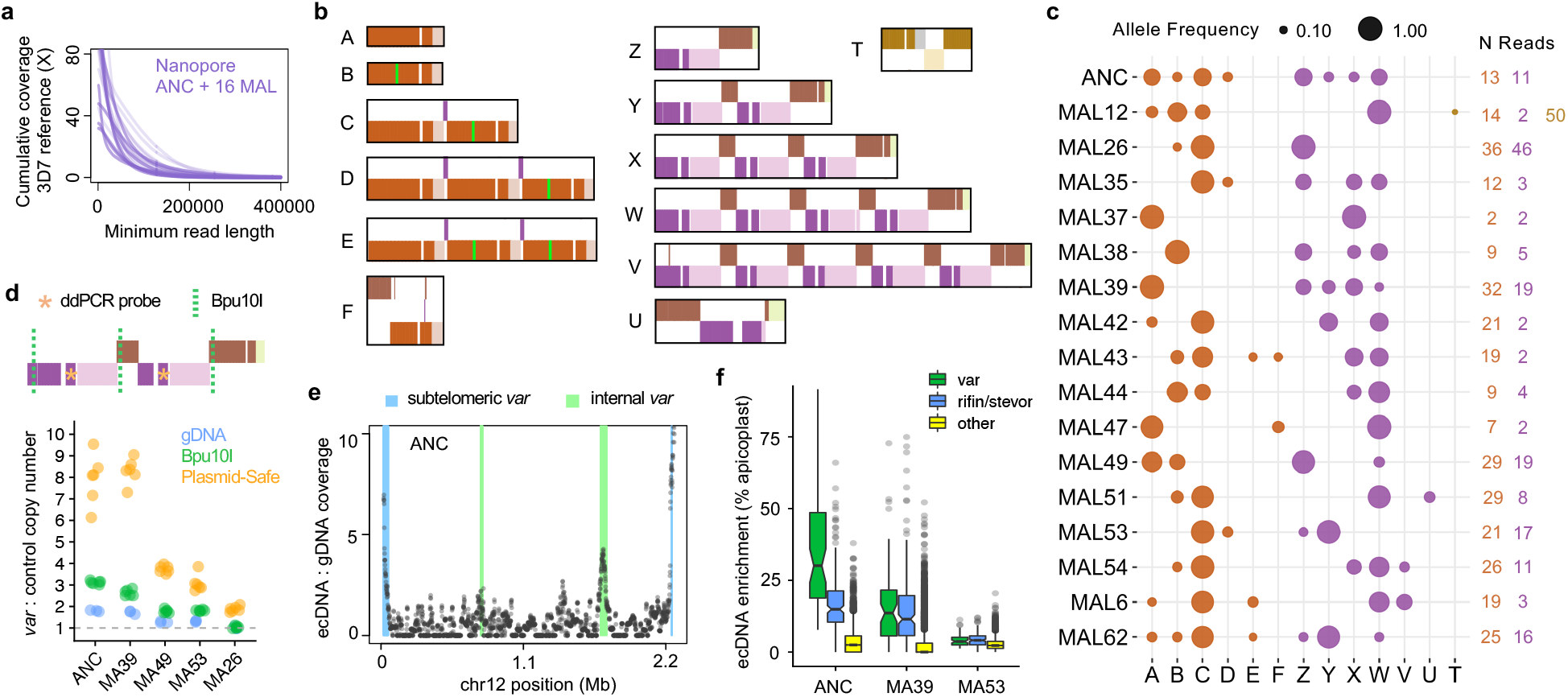
Structural variation at *var* loci maintained on extrachromosomal, circular DNA (ecDNA). **a**, Nanopore read lengths. **b**, Alleles from single reads mapping to the second internal *var* locus on chr12. **c**, Allele frequencies in independent clones, scored from reads spanning local duplications. **d**, Elevated copy number of PF3D7_1240400. Bpu10I is a control for DNA fragmentation. **e**, Nanopore enrichment of ecDNA over genomic DNA on chr12. **f**, Nanopore enrichment of ecDNA genome-wide.

The distribution of chr12 alleles across MAL (**Fig 2C**) was inconsistent with inheritance of a single chromosome from ANC, instead suggesting polyploidy at this locus. Most MAL contained multiple alleles, many of which were shared across MAL and present in ANC. Overall, each MAL appeared to inherit a unique subset of the many alleles present in ANC. Some alleles detected at low frequency in MAL were not observed in ANC reads. Nonetheless, allele F was detectable by PCR in ANC and other lines (**EDF 2B**), suggesting that alleles unobserved by Nanopore may still occur at low frequency. Furthermore, allele frequencies from the two parts of the locus were uncorrelated across MAL (mean pairwise R^2^_adj_=-0.01, linear models; **Fig 2C**), demonstrating that the two regions are genetically unlinked. These observations are inconsistent with each MAL inheriting one DNA molecule containing the chr12 locus, no matter the total number of *var* duplications. Instead, they are consistent with effective polyploidy at this *var* locus (**Fig 2BC**), which is recapitulated at three more internal *var* loci on chr4 and chr7 (**EDF3**).

In light of these patterns, we hypothesized that common *var* polymorphisms were not generated *de novo* during the MA experiment, but instead maintained on extrachromosomal DNA (ecDNA) inherited from ANC with stochastic loss (**Fig 2C**). ecDNA is ubiquitous in eukaryotes^12^ and has been reported in one *P. falciparum* study to date, where it was implicated in selective amplification of a drug-resistance gene^13^. We performed molecular experiments to quantify *var* polymorphism and test the ecDNA hypothesis, again focusing on the most polymorphic internal locus on chr12. First, we used Southern blot to confirm that DNA fragments containing one or two copies of *var* PF3D7_1240300 (**Fig 2B,** orange) were simultaneously present in clonal MAL (EDF 2C). Next, using droplet digital PCR (ddPCR), we found that DNA molecules containing the *var* gene Pf3D7_1240400 (**Fig 2B,** purple) were up to 1.8X more abundant than a control gene on the same chromosome (**Fig 2D**; all p≤0.003 except MA26 p=0.93, t-tests). The copy number of DNA fragments containing this *var* gene within each MAL was strongly correlated with the number of alleles detected with Nanopore (R^2^=0.987, p=0.0004, linear model). Finally, we treated DNA with Plasmid-Safe exonuclease, which efficiently degrades linear but not circular DNA. Plasmid-Safe treatment prior to ddPCR elevated the copy number of Pf3D7_1240400 relative to the control gene in all tested samples (**Fig 2D,** all p<9.3×10^−5^, t-tests; **Supplementary Table 2**), suggesting that extrachromosomal alleles of this *var* locus are maintained on circular DNA.

To obtain a genome-wide estimate of ecDNA, we performed Nanopore sequencing of ANC DNA digested with Plasmid-Safe and aligned the reads to the 3D7 reference genome. After normalization to genomic DNA coverage, ecDNA coverage displayed clear intra-chromosomal peaks at the four internal loci with many common alleles (**Fig 2E,** chr12; **EDF 4**, others). Genome-wide, *var* genes were strongly enriched for ecDNA despite variation in total ecDNA levels across clones (**Fig 2F**; all p<2.2×10^−16^, KS-tests). Besides *var*, the largest gene categories enriched for ecDNA were rifin, STEVOR, and “conserved *Plasmodium* protein unknown function.”

All *P. falciparum* telomeres and subtelomeres were also highly enriched for ecDNA, except on chr14, which contains no *var* (**Fig 2E; EDF 4**). Visual inspection of Nanopore reads and PacBio contigs from subtelomeric *var* loci failed to identify any common polymorphisms shared across lines. Nonetheless, we detected 17 instances of subtelomeric recombination private to individual lines, including 13 found only on single reads (**Supplementary Table 3**). In 7 of these events, sequence from one telomere was copied onto another, creating a chimeric *var* in the recipient locus without altering the donor. Two similar events, along with a duplication of four rifins on chr10, were fixed or nearly fixed in ANC and all MAL and likely occurred prior to the MA experiment. Within individual MAL, we observed six additional instances of telomere replacement that did not interrupt coding genes, including a fixed, reciprocal exchange of non-coding DNA between the second telomeres of chr2 and chr3 in MAL42. Four single reads displayed smaller recombination events, in which 3-7.5 kb of a *var* gene from one subtelomere was overwritten with sequence from another. Together, these rare polymorphisms are consistent with recombination events that occurred during MAL expansion for DNA extraction. We used Luria-Delbruck fluctuation analysis to estimate a *de novo* subtelomeric recombination rate of 6.67×10^−4^ per genome per cell division, which is 3.8-fold lower than a previous estimate from Illumina reads^9^.

*P. falciparum var* loci are known to be enriched for sequence motifs predicted to form G4-quadruplex secondary structures^14^, which are associated with *var* recombination^15^ and replication stalling^16^ that might potentiate ecDNA formation. We noticed that three hypervariable internal *var* loci on chr12, chr7, and chr4 share at least one conserved copy (≥85% identity) of a 7-kb sequence containing six predicted motifs^17^ for G4-quadruplexes. In PlasmidSafe-treated DNA, we also observed many Nanopore reads with central breakpoints for large, inverted duplications (**EDF 5**), which have been implicated in ecDNA formation^18,19^. As expected from Nanopore sequencing of true inverted duplications^20^, the inverse, repeated sequence on the second half of these reads was degraded in sequence quality (**EDF 5C**). Although many questions remain regarding the full mechanism of ecDNA formation in *P. falciparum*, these observations provide intriguing hints about the roles of repetitive DNA and secondary structure.

ecDNA provides an intuitive mechanism for the maintenance of *var* diversity through population bottlenecks, such as mosquito bites that transmit 1-25 *P. falciparum* cells to humans^21^. ecDNA is also consistent with previous observations of gene exchange among *P. falciparum* cells via exosomal vesicles^22^. Nonetheless, since parasite populations express only one *var* gene at a time^23^, most *var* alleles on ecDNA are unlikely to be immediately functional. Instead, we propose that the production and maintenance of ecDNA enables rapid diversification of *var* gene sequence. This function is thought to be under strong selection in response to the host immune system and relevant to vaccine efficacy^2–4,24^. Future clinical work could assess whether *var* diversification through ecDNA impacts parasite evasion of host immune responses and malaria severity. More broadly, these findings add to growing evidence from yeast and cancers implicating ecDNA as a mechanism of rapid adaptation^12,25^.

## Supporting information

all-supplemental-materials

## Methods

### Parasite culture and DNA extraction

*P. falciparum* mutation accumulation lines (MAL) were generated as previously described^11^. Briefly, 31 independent MAL were cloned from an isogenic population of the 3D7 reference strain and propagated in red cell culture. Clonal dilution of MAL was performed every 10.5 parasite cycles to reach a theoretical concentration of 0.25 parasites per well. MAL were cryopreserved after 114-267 days of culture, including 11-25 single-cell bottlenecks.

To generate DNA for PacBio sequencing, cryopreserved aliquots of MAL were cultured in 182 cm^2^ flasks at 4% hematocrit. When parasites reached 10% parasitemia, red blood cell pellets were lysed with saponin at a final concentration of 0.01%. Parasite pellets were washed with 1X PBS and used as input for the Genomic-tip DNA extraction kit (Qiagen). DNA was further cleaned with the PowerClean Pro kit (MoBio) and concentrated by ethanol precipitation.

To generate high-molecular-weight (HMW) DNA for Nanopore sequencing, cryopreserved aliquots of MAL were cultured in 10 mL and 40 mL plates at 2% hematocrit, as previously described^26^. When 80-120 mL of culture reached 4-10% parasitemia at schizont stage, red blood cell pellets were lysed with saponin at a final concentration of 0.014%. Parasite pellets were washed in PBS and suspended in 379 μL extraction buffer (0.1M Tris-HCl pH 8.0, 0.1M NaCl, 20 mM EDTA) with 10 μL of 20 mg/mL Proteinase K (Thermo Fisher Scientific), 10 μL SDS (10% w/v), and 2 μL of 10 mg/mL RNAse A (Millipore Sigma). Tubes were incubated at 55°C for 2-4 hr and gently inverted every 30–60 min to minimize DNA shearing, as previously described^27^. DNA was purified from the lysates with an equal volume of 25:24:1 v/v phenol chloroform isoamyl alcohol (Thermo Fisher Scientific) in a 2 mL light phase lock gel tube (Quantabio), followed by an equal volume of chloroform (Millipore Sigma). HMW DNA in the aqueous layer was poured into a fresh tube and precipitated by adding 0.1 volume 3M sodium acetate and 2-2.5 volumes of cold 100% ethanol. A wide-bore tip was used to transfer visible strings of DNA to a low-retention tube, where it was washed with 70% ethanol and partially air-dried. HMW DNA was resuspended by adding 100-150 uL of 10 mM Tris and incubating at 55°C for up to one hour. To achieve homogenous concentration in the viscous and highly concentrated sample of HMW DNA, samples were gently sheared 1-5 times with a 26G blunt-end needle and incubated for at least 2 weeks at 4°C before proceeding to library preparation. DNA was quantified with Qubit (ThermoFisher) using the dsDNA BR kit.

### PacBio sequencing and genome assembly

SMRTbell libraries were prepared by the Genomics Core at Washington State University and sequenced on a PacBio RS II sequencer. Genomes were assembled *de novo* with HGAP3^28^ using the longest 25X of reads (equivalent to a minimum read length of 16.5-24.7 kb), with additional reads used for polishing. Lowercase (low-quality) bases were removed from the ends of assembled contigs, and contigs containing ≥1000 basepairs with ≥80% identity to the human genome (GRCh37) were identified with BLAST and removed. Remaining contigs were aligned to the *P. falciparum* reference genome (3D7) using minimap2^29^. Dot plots were generated using dotPlotly^30^.

### Nanopore sequencing

Nanopore libraries were prepared from 3-5 μg gDNA with the ONT Ligation Sequencing Kit (SQK-LSK109), with the following modifications to the official protocol to optimize recovery of ultra-long (>100 kb) reads. After the end-prep/repair step, size selection was performed with Short Read Eliminator (SRE) buffer (Circulomics) instead of magnetic beads. After adapter ligation, DNA was isolated by centrifuging the sample at 10,000 × *g* for 30 minutes without the addition of any reagents. This DNA pellet was washed twice with 100 μL SFB or LFB from the ligation sequencing kit and resuspended into 30 μL 10mM Tris pH 8.0. After library preparation, the SRE buffer was used for a final round of size selection, with two washes of SFB or LFB instead of ethanol.

Approximately 350 ng of prepared library was loaded onto the Nanopore flow cell for each sequencing run. To improve throughput, flow cells were flushed every 8-16 hours with the ONT Flow Cell Wash Kit (EXP-WSH003) and reloaded with fresh library. Raw Nanopore data were basecalled with Guppy v3.2.4, using the high-accuracy caller (option: -c dna_r0.4.1_450bps_hac.cfg)

To generate Nanopore reads from ecDNA, 15-20 μg of genomic DNA (gDNA) was digested with ∼95 U Plasmid-Safe ATP-Dependent DNase (Lucigen), 25 mM ATP solution, and Plasmid-Safe 10X buffer for 16 hours at 37°C. The remaining DNA was purified via ammonium acetate precipitation. This treatment eliminated 95-99% of the input DNA, as measured by Qubit (ThermoFisher) and confirmed with gel electrophoresis. Putative ecDNA was gently sheared 10× with a 26G blunt-end needle, and library preparation and sequencing were performed with the standard LSK109 kit protocol (Oxford Nanopore). Library preparation inputs (25-100 ng) and yields (10-25 ng) were far below Nanopore minimum recommendations due to small quantities of DNA remaining after Plasmid-Safe digestion.

### Visual genotyping of *var* loci

Thirty-five *var* loci were defined as 10-110 kb regions of the 3D7 reference genome containing at least one 1 *var* gene or pseudogene and a few surrounding genes. 3D7 sequences at these loci were divided into 500-bp segments and compared with BLAST against PacBio contigs longer than 5 kb and Nanopore reads longer than 30 kb. To visualize the BLAST output, each contig and read was assigned to at most one *var* locus using minimap2 alignment to 3D7. For each locus, a custom Shiny app (available at https://github.com/emily-ebel/varSV) was used to visualize the orientation of the 500-bp reference segments on each individual contig or read. In this visualization, wild-type (3D7) reads or contigs appear as a continuous diagonal line of blocks, similar to a dot plot. Duplications are visible as blocks that deviate from the diagonal line, while deletions appear as truncated or missing genes. Recombination and translocation events are detectable by gaps or truncations in the series of blocks on the diagonal line (due to lack of homology between the read and assigned locus). For putative recombination events >2 kb, we used BLAST and minimap2 to search for donor sequence from elsewhere in the 3D7 reference genome. We did not investigate common, smaller runs of missing segments (<2 kb) after determining that the vast majority were explained by very simple repeats or low-quality sequence. Reads were generally considered wild-type if they contained at least two genes from the assigned locus without structural variation >2 kb. When the same structural variant was detected on >1 read in a sample, we limited the definition of wild-type to reads that were long enough to span the entire variant. Large inverted duplications (‘triangle reads’) were not included in visual allele counts but were quantified from PAF alignments using a custom R script.

### Mutation rate calculation

Mutation rate was estimated using Luria-Delbruck fluctuation analysis via maximum likelihood on the FALCOR web server^31^. This approach assumes exponential growth from a single cell to the large population used for DNA extraction, analogous to a plating experiment on selective media. Rates were calculated for all *var* loci excluding the four hypervariable internal loci on chr12, chr4, and chr7. Each MAL was treated as an independent replicate. The total number of genotyped Nanopore reads was considered the number of viable cells. Reads with the least abundant genotype were considered mutants; i.e., each MAL was assumed to have inherited its most common allele from ANC, which then mutated to produce additional alleles. Deletions on single reads were excluded from the calculation, since they are the most common form of Nanopore sequencing error^32^. Fixed variants within MAL were also excluded, since they are unlikely to have occurred during the final population expansion.

### Coverage analysis

Nanopore reads ≥30 kb from untreated gDNA and ≥10 kb from Plasmid-Safe-treated ecDNA were aligned to the 3D7 reference genome using minimap2. Alignments with MAPQ<10, including multiply mapped reads with MAPQ=0, were removed using samtools^33^ view -q 10. Coverage was calculated in 1-kb windows using bedtoolscoverage and normalized to 1 via division by the genome-wide mean, excluding the circular apicoplast. For analysis by gene category, coverage was calculated in windows corresponding to Ensembl gene annotations (ASM276v2.54) and normalized by the ecDNA:gDNA ratio for the apicoplast, which ranged from 12.0-35.9.

### PCR and Southern Blot

PCR primers were designed to amplify over novel junctions and corresponding reference junctions in the second internal *var* locus on chr12 (**EDF 2A**; **EDT 1**). PCR was performed using Phusion High-Fidelity DNA Polymerase (NEB) according to manufacturer instructions. PCR products were visualized on 1% agarose gels with a 1 kb plus ladder (NEB).

For a Southern Blot probe, primers were designed to amplify a 522-bp segment of PF3D7_1240300 (**EDF 2C; EDT 1**). Amplification was performed on MA53 DNA using Phusion High-Fidelity DNA Polymerase (NEB). The PCR product was purified using the QIAquick PCR Purification Kit (Qiagen) and labeled using the North2South Biotin Random Prime Labeling Kit (ThermoFisher), with the addition of 1.5 uL Glycoblue Precipitant (ThermoFisher). Restriction digests of *P. falciparum* DNA were performed using 10 U SacI-HF (NEB), 5 U StuI (NEB), CutSmart buffer (NEB), and ∼1 ug DNA for 16 hours at 37°C. Digested DNA was run on a gel containing 0.4% UltraPure agarose (Invitrogen) for 7 hours with a current of 3V/cm. TAE buffer was replaced every 2 hours to keep bands crisp. GeneRuler High Range DNA Ladder (ThermoScientific) was used to quantify DNA migration. Gel processing and blotting was performed with the Amersham ECL Direct Nucleic Acid Labeling and Detection Systems Kit (Cytiva) and Amersham Hybond-N+ Nylon Membrane (Cytiva). Probe hybridization was performed using the North2South Chemiluminescent Hybridization and Detection Kit (ThermoScientific). The final blot was visualized with Odessey-XF Imaging System (Li-Cor).

### ddPCR

ddPCR was performed using the QX200 Droplet Digital PCR system (Bio-Rad) and ddPCR Supermix for probes (Bio-Rad). Briefly, primers were designed (**EDT 1**) to amplify a 120-bp fragment of the *var* gene PF3D7_1240400 and a 130-bp fragment of the control gene PF3D7_1212500, which encodes glycerol-3-phosphate 1-O-acyltransferase. Oligonucleotide probes for these amplicons (**EDT 1**) were labeled with HEX/ZEN/IBFQ and FAM/ZEN/IBFG reporter fluorophores (IDT), respectively. For Plasmid-Safe treatments, 400 ng of DNA was incubated with 5 U Plasmid-Safe ATP-Dependent DNase (Lucigen), 25 mM ATP solution, and Plasmid-Safe buffer for 16 hours at 37°C. For restriction enzyme treatments, 200 ng of DNA was incubated with 7.5 U Bpu10I (NEB) in r3.1 buffer (ThermoFisher) for 4 hours at 37°C. ddPCR was performed in five replicates per sample on DNA diluted to 0.05 ng per well. The copy number of *var* and control fragments in each well was calculated using QuantaSoft software (Bio-Rad).

## Legends for Extended Data Figures and Supplementary Tables

**Extended Data Fig. 1: Structural variation among assembled PacBio contigs. a**, Genome-wide dot plots. Insets are internal *var* with >1 contig. **b**, Summary of loci with >1 contig across assemblies. **c**, Visualization of indel polymorphism across contigs at two internal *var* loci on chr4.

**Extended Data Fig. 2: Molecular confirmation of long-read polymorphism. a**, PCR of breakpoints observed in contigs that map to the second internal *var* locus on chr12. Colored bands represent amplicons. Asterisks indicate the amplicons expected in each sample, based on assembled PacBio contigs. In the left diagram, the pink amplicon is expected to be 562 bp in the reference allele (e.g. allele A) and 667 bp with the gene conversion from PF3D7_0700100 (e.g. allele B). These data confirm the existence of breakpoints detected with PacBio in ANC and MA53 but undetected in MA39 and MA47. **b**, PCR detection of allele F. Asterisks indicate the amplicons expected in each sample, based on Nanopore reads. **c**, Southern blot of copy number variation in PF3D7_1240300. Teal asterisks mark the bands expected in each sample, based on Nanopore reads.

**Extended Data Fig. 3: Structural polymorphism across Nanopore reads mapping to internal *var* loci from all clonal MAL**.

**Extended Data Fig. 4: Genome-wide coverage of extrachromosomal, circular DNA relative to genomic DNA**.

**Extended Data Fig. 5: Large inverted duplications on ecDNA reads. a**, Example read from MA54 containing a large inverted duplication. **b**, Plasmid-Safe-treated DNA is strongly enriched for “triangle reads”. **c**, Signal degradation consistent with single-strand annealing after passing through Nanopore.

**Supplementary Table 1. Summary of PacBio assemblies including PAF alignments to 3D7**.

**Supplementary Table 2. ddPCR count data from QuantaSoft**.

**Supplementary Table 3. Structural variation on Nanopore reads assigned to non-hypervariable *var* loci**. The three events fixed in ANC and all MAL are considered wild-type.

**Extended Data Table 1.**
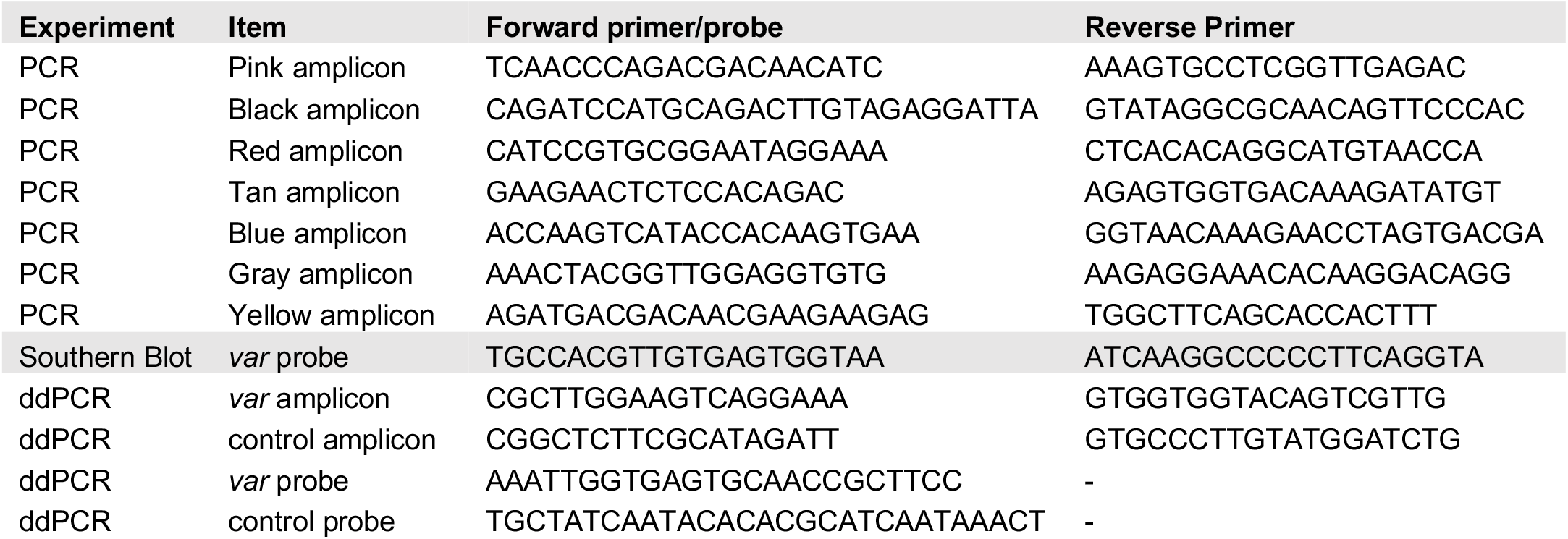
Probe and primer sequences. Colors refer to diagrams in EDF 2.

## Data Availability Statement

Read data and genome assemblies generated by this study are deposited at NCBI SRA and GenBank under NCBI BioProject: PRJNA894225.

## Code Availability Statement

Code used to identify structural variants is available at https://github.com/emily-ebel/varSV.

## Acknowledgements

We thank Ryan Taylor, Jennifer Guler, Shiwei Liu, Xue Li, and Thomas Braukmann for helpful discussion; Melanie Espiritu, Nana Ansuah Petersen, and Mark Wildung for laboratory assistance; Stefan Oliver, Katja Pekrun, and Asuka Eguchi for loaning equipment; and Alison Feder, Grant Kinsler, Kerry Geiler-Samerotte, and future reviewers for their helpful comments on the manuscript. This work was supported by a seed grant from the Stanford Center for Computational, Evolutionary and Human Genomics (CEHG) to ERE; a CEHG predoctoral fellowship to ERE; NIH grant F32GM135998 to BYK; NIH grant 1DP2HL13718601 to ESE; NIH grant R37AI048071 to TJCA; and NIH grant R35GM118165 to DAP. ERE is an NSF Postdoctoral Research Fellow in Biology (#2109851). ESE is a Tashia and John Morgridge Endowed Faculty Scholar in Pediatric Translational Medicine through the Stanford Maternal Child Health Research Institute. ESE and DAP are Chan Zuckerberg Biohub Investigators.

## Author contributions

ERE, DAP, and TJCA conceptualized the study. MMW, ERE, and ESE cultured the parasites. ERE and MMW isolated DNA. BYK performed Nanopore sequencing. ERE performed the other experiments. ERE and BYK analyzed sequencing data. ESE, TJCA, and DAP contributed resources and supervision. ERE and DAP wrote the manuscript with input from all authors.

## References

1. World Health Organization. World malaria report 2020: 20 years of global progress and challenges. (World Health Organization, 2020).

2. Bull, P. C. et al. Parasite antigens on the infected red cell surface are targets for naturally acquired immunity to malaria. Nat. Med. 4, 358–360 (1998).

3. Langhorne, J., Ndungu, F. M., Sponaas, A.-M. & Marsh, K. Immunity to malaria: more questions than answers. Nat. Immunol. 9, 725–732 (2008).

4. Saito, F. et al. Immune evasion of Plasmodium falciparum by RIFIN via inhibitory receptors. Nature 552, 101–105 (2017).

5. Su, X. Z. et al. The large diverse gene family var encodes proteins involved in cytoadherence and antigenic variation of Plasmodium falciparum-infected erythrocytes. Cell 82, 89–100 (1995).

6. Rask, T. S., Hansen, D. A., Theander, T. G., Pedersen, A. G. & Lavstsen, T. Plasmodium falciparum Erythrocyte Membrane Protein 1 Diversity in Seven Genomes – Divide and Conquer. PLOS Comput. Biol. 6, e1000933 (2010).

7. Chen, D. S. et al. A Molecular Epidemiological Study of var Gene Diversity to Characterize the Reservoir of Plasmodium falciparum in Humans in Africa. PLOS ONE 6, e16629 (2011).

8. Otto, T. D. et al. Evolutionary analysis of the most polymorphic gene family in falciparum malaria. Wellcome Open Res. 4, 193 (2019).

9. Claessens, A. et al. Generation of Antigenic Diversity in Plasmodium falciparum by Structured Rearrangement of Var Genes During Mitosis. PLOS Genet. 10, e1004812 (2014).

10. Otto, T. D. et al. Long read assemblies of geographically dispersed Plasmodium falciparum isolates reveal highly structured subtelomeres. Wellcome Open Res. 3, 52 (2018).

11. McDew-White, M. et al. Mode and Tempo of Microsatellite Length Change in a Malaria Parasite Mutation Accumulation Experiment. Genome Biol. Evol. 11, 1971–1985 (2019).

12. Noer, J. B., Hørsdal, O. K., Xiang, X., Luo, Y. & Regenberg, B. Extrachromosomal circular DNA in cancer: history, current knowledge, and methods. Trends Genet. TIG 38, 766–781 (2022).

13. McDaniels, J. M. et al. Extrachromosomal DNA amplicons in antimalarial-resistant Plasmodium falciparum. Mol. Microbiol. 115, 574–590 (2021).

14. Smargiasso, N. et al. Putative DNA G-quadruplex formation within the promoters of Plasmodium falciparum var genes. BMC Genomics 10, 362 (2009).

15. Stanton, A., Harris, L. M., Graham, G. & Merrick, C. J. Recombination events among virulence genes in malaria parasites are associated with G-quadruplex-forming DNA motifs. BMC Genomics 17, 859 (2016).

16. Bryan, T. M. Mechanisms of DNA Replication and Repair: Insights from the Study of G-Quadruplexes. Molecules 24, 3439 (2019).

17. Kikin, O., D’Antonio, L. & Bagga, P. S. QGRS Mapper: a web-based server for predicting G-quadruplexes in nucleotide sequences. Nucleic Acids Res. 34, W676–W682 (2006).

18. Nonet, G. H., Carroll, S. M., DeRose, M. L. & Wahl, G. M. Molecular Dissection of an Extrachromosomal Amplicon Reveals a Circular Structure Consisting of an Imperfect Inverted Duplication. Genomics 15, 543–558 (1993).

19. Grondin, K., Roy, G. & Ouellette, M. Formation of extrachromosomal circular amplicons with direct or inverted duplications in drug-resistant Leishmania tarentolae. Mol. Cell. Biol. (1996) doi:10.1128/MCB.16.7.3587.

20. Spealman, P., Burrell, J. & Gresham, D. Inverted duplicate DNA sequences increase translocation rates through sequencing nanopores resulting in reduced base calling accuracy. Nucleic Acids Res. 48, 4940–4945 (2020).

21. Beier, J. C., Davis, J. R., Vaughan, J. A., Noden, B. H. & Beier, M. S. Quantitation of Plasmodium falciparum sporozoites transmitted in vitro by experimentally infected Anopheles gambiae and Anopheles stephensi. Am. J. Trop. Med. Hyg. 44, 564–570 (1991).

22. Regev-Rudzki, N. et al. Cell-Cell Communication between Malaria-Infected Red Blood Cells via Exosome-like Vesicles. Cell 153, 1120–1133 (2013).

23. Chookajorn, T., Ponsuwanna, P. & Cui, L. Mutually exclusive var gene expression in the malaria parasite: multiple layers of regulation. Trends Parasitol. 24, 455–461 (2008).

24. Raghavan, M. et al. Proteome-wide antigenic profiling in Ugandan cohorts identifies associations between age, exposure intensity, and responses to repeat-containing antigens in Plasmodium falciparum. http://biorxiv.org/lookup/doi/10.1101/2022.06.24.497532 (2022) xdoi:10.1101/2022.06.24.497532.

25. Demeke, M. M., Foulquié-Moreno, M. R., Dumortier, F. & Thevelein, J. M. Rapid Evolution of Recombinant Saccharomyces cerevisiae for Xylose Fermentation through Formation of Extra-chromosomal Circular DNA. PLOS Genet. 11, e1005010 (2015).

## Methods References

26. Ebel, E. R., Kuypers, F. A., Lin, C., Petrov, D. A. & Egan, E. S. Common host variation drives malaria parasite fitness in healthy human red cells. eLife 10, e69808 (2021).

27. Kim, B. Y. et al. Highly contiguous assemblies of 101 drosophilid genomes. eLife 10, e66405 (2021).

28. Chin, C.-S. et al. Nonhybrid, finished microbial genome assemblies from long-read SMRT sequencing data. Nat. Methods 10, 563–569 (2013).

29. Li, H. Minimap2: pairwise alignment for nucleotide sequences. Bioinformatics 34, 3094–3100 (2018).

30. Poorten, T. dotPlotly. (2022).

31. Hall, B. M., Ma, C.-X., Liang, P. & Singh, K. K. Fluctuation analysis CalculatOR: a web tool for the determination of mutation rate using Luria-Delbruck fluctuation analysis. Bioinforma. Oxf. Engl. 25, 1564–1565 (2009).

32. Delahaye, C. & Nicolas, J. Sequencing DNA with nanopores: Troubles and biases. PLOS ONE 16, e0257521 (2021).

33. Li, H. et al. The Sequence Alignment/Map format and SAMtools. Bioinformatics 25, 2078–2079 (2009).

34. Quinlan, A. R. & Hall, I. M. BEDTools: a flexible suite of utilities for comparing genomic features. Bioinformatics 26, 841–842 (2010).

