## Supplementary figures and images for "Antigenic diversity in malaria parasites is maintained on extrachromosomal DNA"

### EDF1-PacBio.pdf

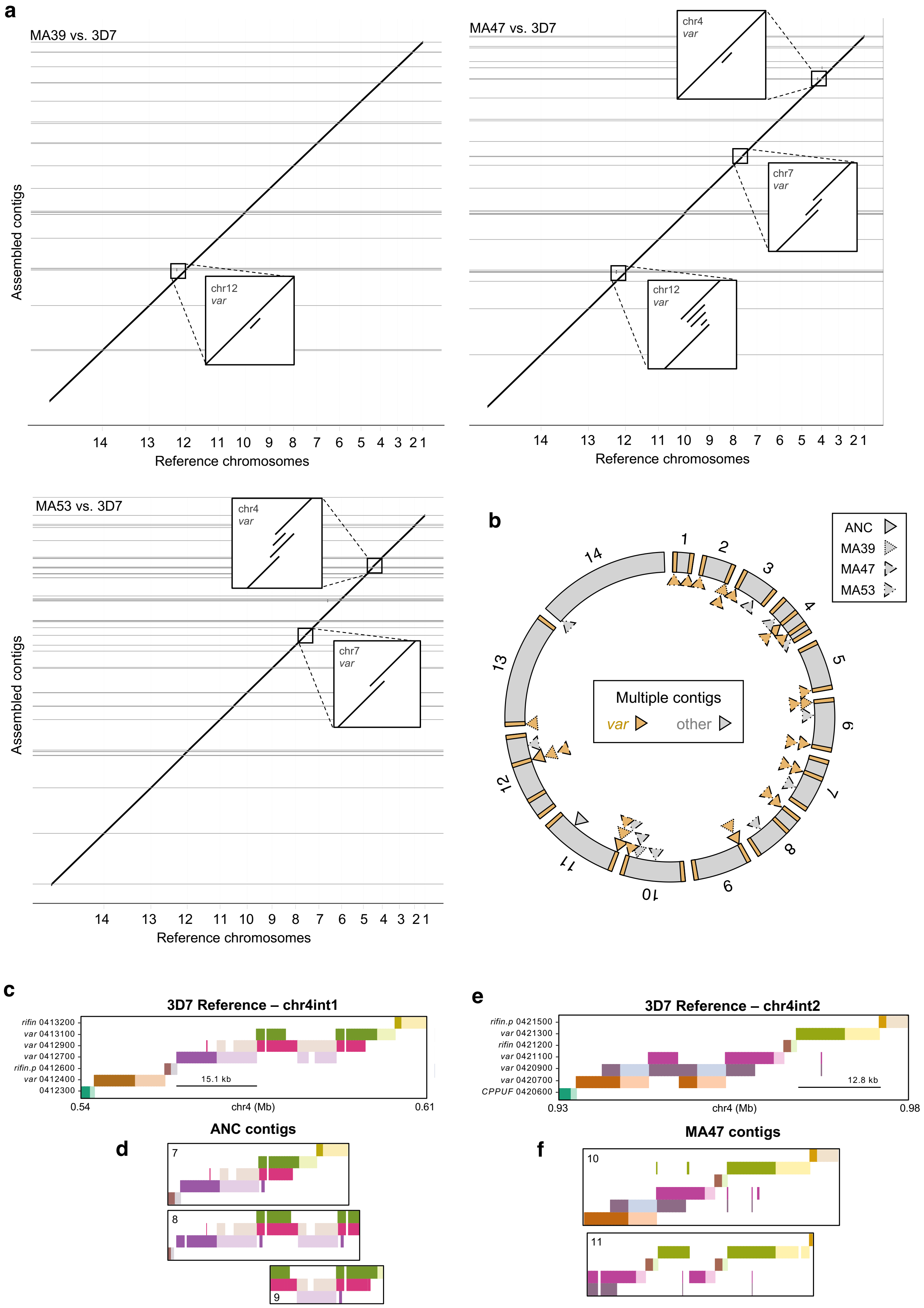

### EDF2-PCR-Sblot.pdf

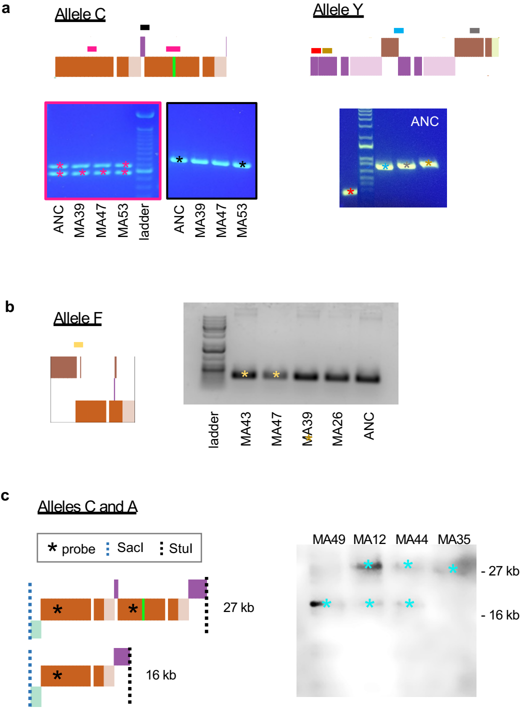

### EDF3-Nanopore-chr7-chr4.pdf

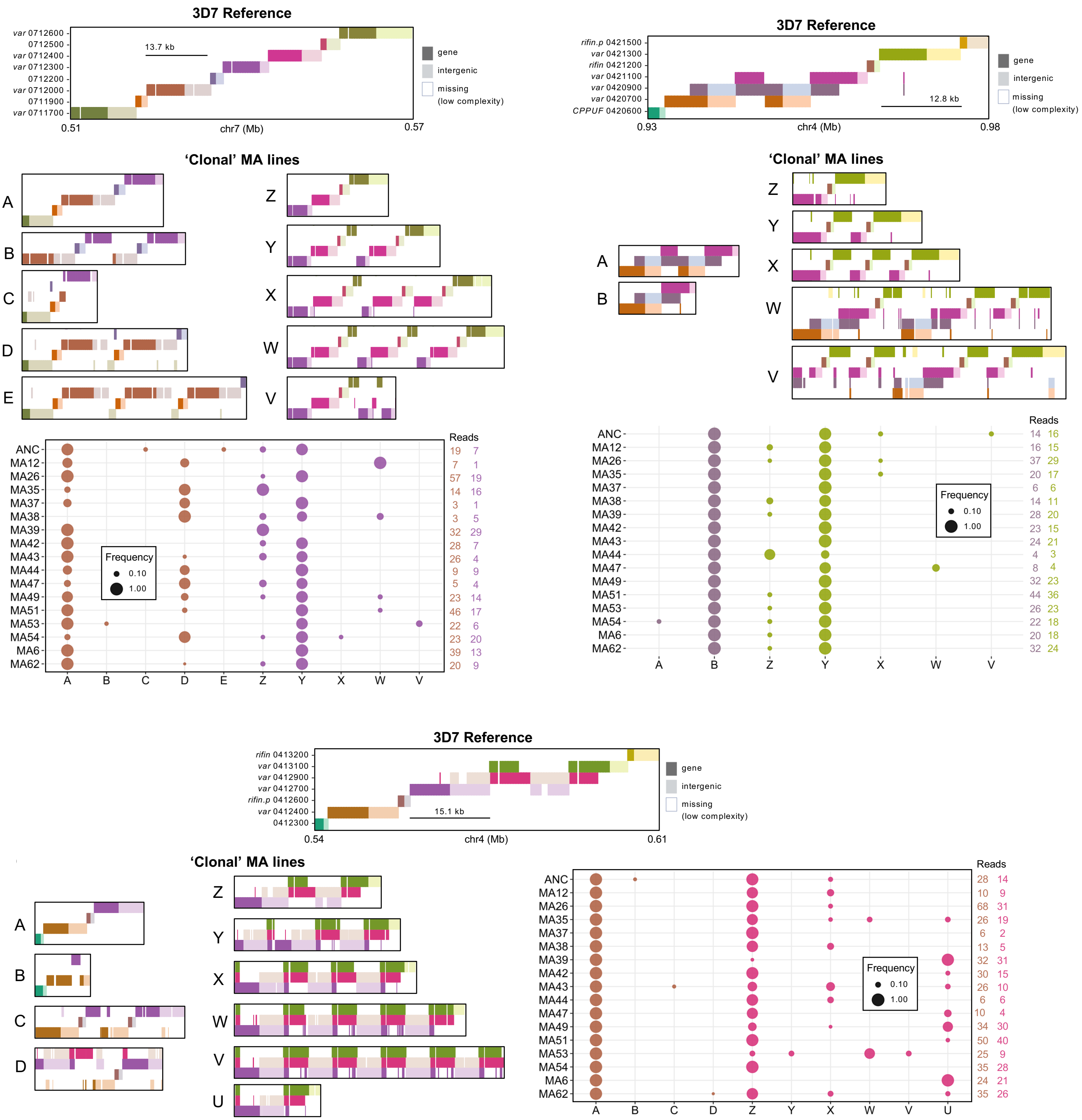

### EDF4-ecDNA-coverage.pdf

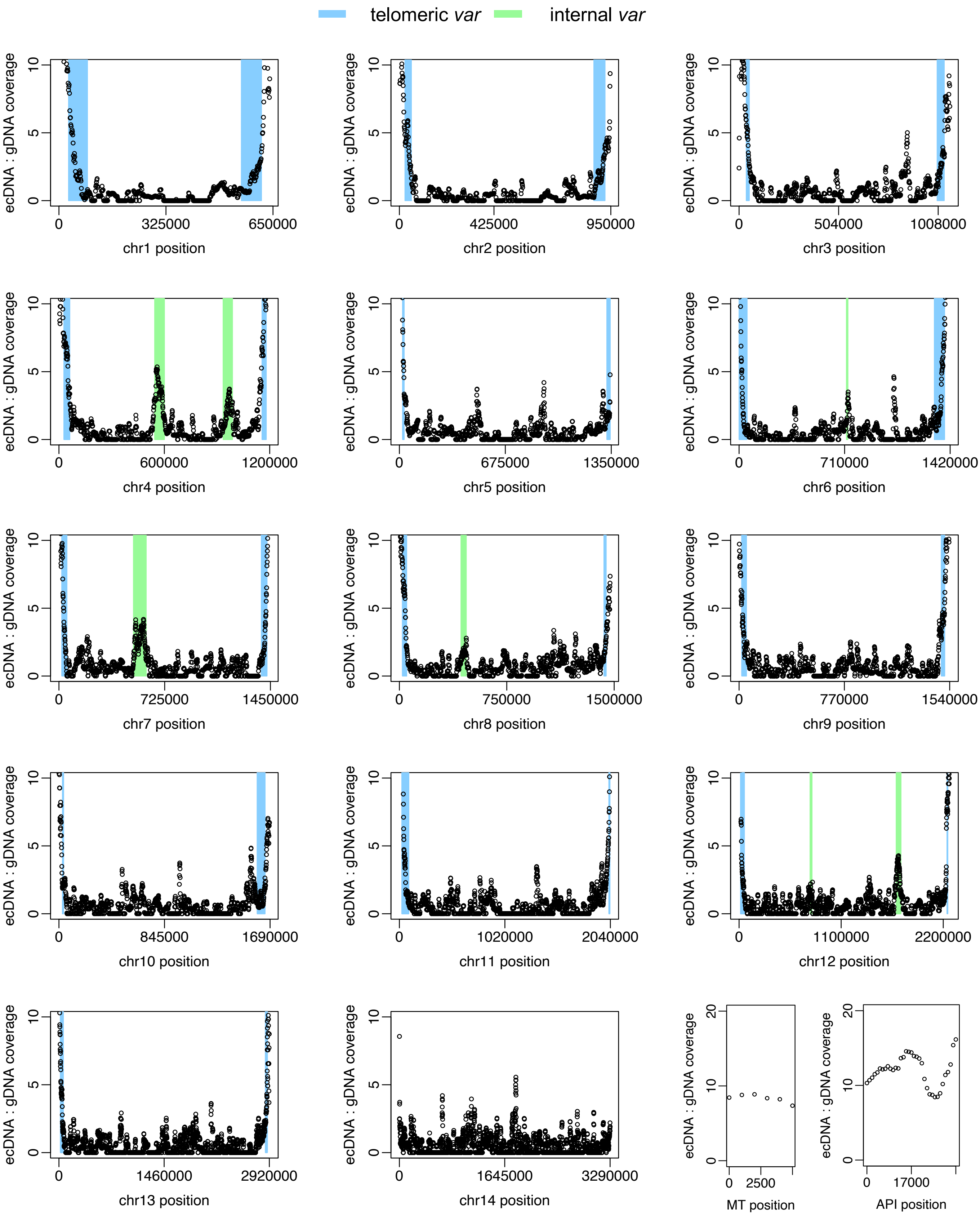

### EDF5-Triangles.pdf

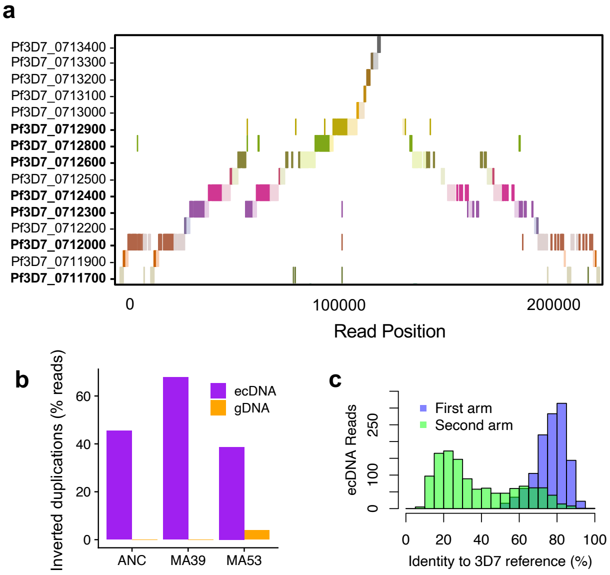
